# Apurinic/apyrimidinic endodeoxyribonuclease 2 (*APEX2*/APE2) is required for efficient expression of *TERT* in human embryonic stem cells

**DOI:** 10.1101/2024.09.23.614488

**Authors:** Josh L. Stern, Lindsay F. Rizzardi, Natalie R. Gassman

## Abstract

Human stem cells rely on enhanced DNA repair mechanisms to safeguard their ability to replenish somatic tissues. Telomerase counteracts telomere shortening and is a component of the stem cell DNA repair system that is regulated by ATM and ATR kinases. Here, we report that the DNA repair enzyme APEX2, but not its close paralog APEX1, is required for efficient telomerase reverse transcriptase (*TERT*) gene expression in human embryonic stem cells (hESC) and a melanoma cell line. We also observed that APEX2 knockdown significantly diminished telomerase enzyme activity. While APEX1 is known to regulate certain transcription factors, APEX2 has not been reported to influence gene expression. To gain insight into how APEX2 influences gene expression, we conducted RNA-seq following APEX2 knockdown in hESC. These results indicated that a number of genes, in addition to *TERT*, relied on APEX2 for efficient expression. Genes affected by APEX2 knockdown were significantly enriched for specific repetitive DNA families. These include mammalian-wide interspersed repeats (MIRs) and *Alu* elements. Chromatin immunoprecipitation experiments demonstrated the highest APEX2 binding near MIR sequences in *TERT* intron 2. Surprisingly, binding was low in the *TERT* proximal promoter, a region known to control *TERT* transcription. MIR and other repetitive DNA regions are common sites of DNA damage, suggesting that APEX2 recruitment and repair of *TERT* MIR sequences may play a role in influencing *TERT* expression. This new role for APEX2 in promoting efficient gene expression deepens our understanding of an emerging cancer therapeutic target. Further, as the *TERT* gene plays critical roles in stem cell maintenance, organismal development and aging, as well as in short telomere disorders and cancer, our observations provide insight into new strategies to modulate the expression of this important enzyme.

## INTRODUCTION

Human stem cells express telomerase, a specialized cellular DNA repair system enzyme that counteracts telomere shortening (1–3). The catalytic subunit, encoded by the *TERT* gene, plays critical roles in stem cell maintenance, organismal development and aging and is often dysregulated in short telomere diseases and cancer (4–6). *TERT* mRNA transcription is tightly regulated and is largely restricted to stem cells. The mature telomerase enzyme contains several factors, but control of *TERT* transcripts is currently considered the major on/off switch restricting telomerase expression in humans (7–10).

*TERT* is haploinsufficient, and patients with hypomorphic mutations in telomerase components typically have short telomeres and display a range of premature aging characteristics (11), such as bone marrow failure, liver problems, cancer, and premature death (12–14). Increasing telomere lengths in models of these conditions suggests that enhancing telomerase levels could benefit patients (9, 11, 15–22). In addition, telomere shortening causes stem cells to become dysfunctional in aging tissues and studies suggest that controlling telomerase expression may lead to healthier aging (reviewed in (4)).

Historical challenges have left gaps in our understanding of *TERT* transcriptional control. *TERT* is expressed at low levels in human stem cells (23), making it difficult to study *TERT* at the protein level (24, 25). In addition, 50% decreases in telomerase expression typically have significant biological effects on telomere dynamics in humans (26–28). The regulation of mouse *TERT* (m*TERT*) and human *TERT* (h*TERT*) differ (29, 30) limiting the suitability of murine models for studying the developmental regulation of *TERT* transcription. Although 90% of cancers express telomerase, the mechanisms driving *TERT* expression vary widely and it remains unclear which cancer cells, if any, might be suitable models for normal human stem cells (31–38). Induced pluripotent stem cells (iPSC) have been used to study *TERT*. However, telomerase in these cells is artificially activated and how well they model normal human stem cells is unknown. Contributing to these challenges, validated models to study normal human stem cells have been lacking until recently.

DNA repair in human stem cells is particularly important because they need to replicate numerous times without DNA errors. Such errors, left unrepaired, can negatively impact the function of the tissues the stem cells support. Because of this special importance in stem cells, DNA repair mechanisms are enhanced in this cell type and DNA repair failures in stem cells lead to a range of diseases (39–41). How stem cells maintain genome integrity is an active area of investigation.

Many DNA repair factors regulate gene expression (42, 43). For example, in human cells, apurinic/apyrimidinic endodeoxyribonucleases (*APEX1*) APEX1 is considered the major AP endonuclease but also directly influences transcription through its redox effector (REF-1) domain. Humans also encode a second apurinic/apyrimidinic endodeoxyribonuclease, *APEX2* (a.k.a. APE2). While APEX2 is involved in many aspects of DNA maintenance, including base excision repair, single-strand break repair, microhomology mediated end joining, immunoglobulin class switch recombination and somatic hypermutation (44), prior studies have not identified roles for APEX2 in regulating gene transcription. Of note, in the scientific literature, the plant gene *APX2* (L-ascorbate peroxidase 2) is widely misnamed APEX2 and should not be confused with human APEX2 studied here. Here we report a previously unrecognized role for APEX2 in promoting efficient expression of a subset of genes, including *TERT*, a function that may rely on its ability to resolve DNA damage occurring in repetitive sequences (45–51).

## 2. MATERIALS AND METHODS

### Cell culture

Human pluripotent embryonic stem cells (WA01) were obtained from Wicell Inc., and induced pluripotent stem cells derived from reprogramming CD34^+^ hematopoietic cells (BXS0116) were obtained from American Type Cell Culture Collection. These cells were maintained in feeder-free culture on vitronectin (Gibco, A14700) in Essential 8 Flex media (Gibco, A2858501), supplemented with glutamine analog SG-200 (Cytiva, SH30590.01) and 1% Penicillin-Streptomycin (Cytiva, SV30010). For passaging, cells were dissociated using 0.5 mM EDTA, washed in complete medium, and then reseeded with 10 µM ROCK inhibitor Y-27693 (ApexBio) overnight, followed by replacement with complete media. WA01 cells are derived from the inner cell mass (ICM) of the blastocyst stage of embryonic development and retain their capacity to differentiate into cells of different lineages (52, 53). Transcriptomic and phenotypic analyses under these conditions indicated the expression of markers associated with primed pluripotent cells, which includes Sox2, Oct4 and Nanog but not markers associated with naïve pluripotency (54, 55).

Glioma stem cells (GSC) were grown as previously described (56, 57). Melanoma cells (A101) were maintained as an adherent culture in DMEM media (Cytiva, SH30022.02) supplemented with glutamine analog SG-200 (Cytiva, SH30590.01), 1% sodium pyruvate (Corning, 25-000-CI), and 1% penicillin/streptomycin (Cytiva, SV30010). All cell cultures were maintained at 37 °C and 5% CO_2_ in a humidified atmosphere.

Cells were treated with the following small molecule inhibitors, MEK1/2 inhibitor trametinib (ApexBio, A3018) and PD0325901 (ApexBio, A3013), ERK inhibitor SCH772984 (ApexBio, B5866), EZH2 inhibitor GSK343 (ApexBio, A3449) or the c-Myc:MAX dimerization inhibitor 10058-F4 (ApexBio, A1169).

### RNA extraction and cDNA synthesis

RNA was extracted from cells using RNeasy Mini Kit (Qiagen, 74106) and DNase-digestion was performed during RNA extraction using RNase-free DNase (Qiagen, 79254). cDNA synthesis was primed with random hexamers and oligo dT at final concentrations of 1 µM and 5 µM respectively, heated to 65°C for 5 min and then snap-cooled. The RNA samples were then reverse transcribed (Protoscript II, New England Biolabs, M0368X) in the prescribed buffer with 0.1 M DTT, 500 mM dNTPs, and RNase inhibitor (Promega, N2511) at 25°C for 5 min, 42°C for 60 min and 65° for 20 min. The cDNA samples were then treated with RNase H (New England Biolabs, M0297L) and incubated at 37°C for 30 min and 65°C for 20 min. Quantitative PCR was performed using SYBR Select (ThermoFisher, 4472908) using CFX real-time PCR system (Biorad) and CFX Maestro software was used for quantification. All PCR amplicons were sequenced at least once to ensure the accuracy of product obtained. Primers used for amplification are listed in Supplementary Table 1.

### Cell Lysis and Immunoblots

After removing media from cells in a 6-well plate, 1.5 mL of ice-cold PBS was added, and the cells were scraped into a 1.5 mL tube. Cells were collected by centrifugation at 400 × g for 4 min, PBS was removed, and cells were placed on ice for lysis or stored at −80 °C. Cell pellets were lysed in 10 mM Tris-Cl (pH 8.0), 150 mM sodium chloride, 1% Triton X-100, 1 mM EDTA, with Complete Protease Inhibitor (Fisher, A32963) and phosphatase inhibitors (ApexBio, K1015). Samples were incubated on ice for 20 min, then centrifuged for 20 min at 13,000 × g to remove insoluble material. Lowry protein estimation was conducted to quantify the concentrations of cell lysates with DC™ Protein Assay Reagent S (Bio Rad, 500-0115) and DC Protein Assay Reagent A (Bio Rad, 5000113) according to the manufacturer’s protocol. Absorbance readings at 750 nm using a BioTek Synergy 2 plate reader were used to calculate mg/mL protein values. 50 μg of protein were made up in 1 × NuPage LDS sample buffer (Invitrogen, NP0007) and Invitrogen Novex 10 × Bolt Sample Reducing Agent (Thermo Fisher Scientific, B0009), incubated at 95°C for 7 min before loading equal protein amounts onto Mini-PROTEAN TGX Stain-Free Gels 4–20% Tris-Glycine polyacrylamide gels (Bio Rad, 4568096). Gels were run in 1 x TGS buffer for 60 min at 110 volts and transferred to Trans-Blot® Turbo™ Mini Nitrocellulose membranes (Bio Rad, 1704158) for 5 min using the Trans-Blot® Turbo™ RTA Transfer Kit, Nitrocellulose System (Bio Rad, 170-4270), and Trans-Blot Turbo Transfer Buffer (Bio Rad, 10026938). Membranes were blocked in 5% StartingBlock™ (TBS) Blocking Buffer (Fisher, 37542) with orbital shaking for 30 min at room temperature (RT). Primary antibodies were incubated with blots overnight at 4°C with orbital shaking, followed by antibody removal and washing in 10 mL of TBS-T three times for 5 min at RT with orbital shaking. Primary antibodies (Supplementary Table 2) were detected by adding species-specific horseradish peroxidase conjugated-secondary antibody in 5% nonfat dry milk, incubated with orbital shaking overnight at 4°C followed by washing as for primary antibodies. After removing the last wash buffer, chemiluminescent visualization (SuperSignal™ West Pico PLUS Chemiluminescent Substrate, Thermo Fisher Scientific, 34578) was used. Membranes were visualized on a BioRad ChemiDoc MP. Membranes were then stripped three times in a low-pH stripping buffer (25 mM Glycine, 1% SDS, pH 2.3) for 15 min on an orbital shaker and then washed in 10 mL TBS-T for 5 min on the orbital shaker before the new primary antibody was added.

### Chromatin Immunoprecipitation (ChIP)

Adherent cells in 15 cm plates were washed with PBS, then fixed in 1% formaldehyde in PBS for 10 min at room temperature, and then treated with 125 mM glycine for 2 min to inactivate the formaldehyde. The solution was aspirated, and cells were immediately scraped into 15 mL of ice-cold PBS and spun at 500 x g for 5 min; the PBS was removed, and cells were frozen at -80°C. Cells were then lysed for 10 min on ice in 500 mL of 50 mM Tris-Cl pH 8, 10 mM EDTA, 0.5% SDS, complete protease (Fisher, A32963) and phosphatase inhibitors (ApexBio, K1015). Nuclei were disrupted and chromatin solubilized by sonication in a BioRuptor in 1.5 mL tubes until fragments were between 100 and 300 base pairs. The size of fragmented DNA was assessed by running the purified DNA on a 1% agarose gel and visualizing using BioRad GelDoc. Solubilized chromatin was obtained by centrifuging the sonicated lysate at 16,000 x g for 15 min at 16°C and discarding the pellet. The chromatin was quantified using Nanodrop. 10 mg of chromatin were nutated overnight with antibody in 1 mL IP buffer (16.7 mM Tris-Cl pH 8.1, 1.2 mM EDTA, 167 mM NaCl, 1 % Triton X-100) at 4°C. For precipitation, 20 µL of magnetic beads were washed once in IP buffer, then added to samples, nutated at 4°C for 24 hours. Chromatin bound to magnetic beads was isolated using a magnetic rack and washed with 1 mL of low-salt buffer (20mM Tris-Cl pH8.0, 2 mM EDTA, 150 mM NaCl, 0.1% SDS, 1% Triton X-100), 1 mL of high-salt buffer (20 mM Tris-Cl pH8.0, 2 mM EDTA, 500 mM NaCl, 0.1% SDS, 1% Triton X-100), 1 mL of LiCl wash (10mM Tris-Cl pH8.0, 1 mM EDTA, 250 mM LiCl, 1% deoxycholate, 1% NP40) and then 1 mL of TE (10 mM Tris pH 8, 2 mM EDTA) for 1 min each. The TE was removed and the immunoprecipitates were eluted from the beads by incubating in 120 µL of elution buffer (100 mM NaHCO3 and 1% SDS) for 30 min at RT, mixing gently every 10 min. The supernatant was collected, and crosslinks were reversed with 5 mM NaCl at 65°C for 24 hours. The protein and RNA in the samples were digested for 1 h using 7 µL of 1 M Tris pH 6.5, 3 µL of 500 mM EDTA, 3 µL of proteinase K (20 mg/mL), 0.5 µL of RNase A (ThermoFisher, EN5031) and then purified using phenol:chloroform: isoamyl alcohol extraction followed by ethanol precipitation with 0.5 mg glycogen. Input chromatin was purified alongside the immunoprecipitates. Following ethanol precipitation of the purified chromatin, samples were analyzed by quantitative PCR with SYBR Select (ThermoFisher, 4472908) on a CFX real-time PCR system (Bio Rad) and CFX Maestro software was used for quantification. PCR reaction for the *TERT* promoter included 0.5 µL of 7-deaza GTP (Roche, 14020724) per 10 µL reaction. Primers used for amplification are listed in Supplementary Table 1 and antibodies are listed in Supplementary Table 2.

### Repair Assisted Damage Detection (RADD)

Oxidative DNA damage was evaluated using Repair Assisted Damage Detection (RADD), which detects DNA lesions using a cocktail of DNA repair enzymes. For detection of oxidative lesions (oxRADD), Fapy-DNA glycosylase (FPG, NEB M0240) and Endonuclease IV (Endo IV, NEB M0304) are used to remove the oxidative DNA lesions and process the abasic sites, then the gapped DNA is tagged using a digoxigenin-labeled dUTP inserted with Klenow exo-, which lacks proofreading (58, 59). We assessed DNA lesions within the hESC cells by spin-coating 100,000 cells onto a SuperFrost Plus coverslide (Thermo Fisher 12-550-15). The cells were then fixed using 3.7% formaldehyde for 10 min and washed three times with PBS following fixation. Cells were outlined with a Pap-pen and permeabilization buffer (Biotium 22016) with 0.05% Triton-X (Sigma Aldrich 648466) was used to permeabilize the cells for 10 min at 37 °C. The cells were then washed three times with 1× PBS.

The cells were then incubated with the lesion removal cocktail, 4U of FPG+ 5 U of EndoIV, resuspended in 1× ThermPol buffer (New England BioLabs B9004S) + 200 µg/ml bovine serum albumin (BSA, Jackson Immuno 001-000-162) for 1 h at 37 °C in a hybridization FISH oven (59). Once the incubation period has finished, the gap-filling mixture of Klenow exo-(Thermo Fisher EP0422) and digoxigenin-labeled dUTP (Sigma Aldrich 11093088910) is directly added to the chambers and placed again within the oven at 37 °C for an additional 1 h. The slides are washed three times with 1× PBS and blocked with 2% BSA in PBS for 30 min. After blocking, slides are incubated with primary antibody anti-digoxigenin (1:250 ab420 Abcam) or anti-mouse IgG1 isotype control (1:625 5415 Cell Signaling) for 1 h at RT. IgG1 is used as a negative control in these cells. Once the primary antibody has finished incubation, the cells are washed three times with 1× PBS and incubated with secondary antibody anti-mouse AlexaFluor 546 (1:400 Thermo Fisher) for 1 h at RT. The cells are then incubated with Hoechst solution (1:800, Thermo Fisher) for 15 min at RT and washed with 1× PBS three times.

Cells are then imaged using the Keyence microscope with the 20× objective (NA 0.75). At least 30 cells of each condition for each replicate are taken over 4 different fields. For analysis, the Nikon Elements software created an ROI around the nucleus, and the total intensity for the RADD channel within the nucleus was recorded for at least 100 cells. The mean fluorescent intensity for all the nuclei is graphed in GraphPad Prism, and significance was calculated using a Welch’s t-test for treated versus control.

### RADD-IP-qPCR

DNA lesions within a specific promoter as assessed using a modified oxRADD assay similar to (60). DNA from PARP1-inhibited cells (olaparib, 500 nM, 24h) was extracted and subjected to oxRADD. Briefly, a 100 µl reaction of 10 µg isolated DNA with 1 x ThermPol Buffer, 200 µg/ml BSA, 25 U FPG, and 25 U EndoIV was incubated for 1 h at 37^0^C, forming gapped DNA. Then 20 U of Klenow exo-DNA polymerase and 1.5 µl of 1 mM biotinylated dUTP (Thermo Fisher R0081) were incubated for an additional hour at 37^0^C to fill the gap, tagging the damage site. DNA was then precipitated with sodium acetate (Thermo Fisher AM9740) and ethanol at -80^0^C. The DNA was then washed, resuspended, and fragmented by sonication for 1 min and subjected to streptavidin bead-mediated precipitation (New England Biolabs, S1420S) of biotinylated fragments followed by purification and qPCR detection of the *TERT* promoter. As a negative control, cells were processed without unlabeled dUTP.

### RNA sequencing and data analysis

RNA sequencing was performed by the UAB Genomics Core Lab using 500 ng of RNA as input. Sequencing libraries were made using the NEBNext Ultra II Directional RNA-seq kit (NEB #E7760L) as per the manufacturer’s instructions and using the NEBNext Poly(A) mRNA Magnetic Isolation Module (NEB #E7490). The resulting libraries were analyzed using BioAnalyzer 2100 (Agilent) and quantified using qPCR (Roche). Sequencing was performed on the NovaSeq 6000 (Illumina) with paired-end 100 bp chemistry as per standard protocols. Fastqs were processed using the nf-core/rnaseq (v3.14.0) pipeline (61) with the --gcBias flag, aligning to the GRCh38.p13 reference, and using the Gencode v32 transcript annotations.

DESeq2 (62) was used with default settings to identify differentially expressed genes after filtering out genes with less than 10 reads in all three samples within a group. After multiple testing correction using the Benjamini-Hochberg (BH) method (63), an FDR cutoff of 0.05 was used to identify differential expression. Gene ontology analysis was performed using enrichr (64) package and “GO_Biological_Process_2023”. To determine enrichment of repetitive elements within this set of differentially expressed genes, we used Poly-Enrich(65). We obtained genomic coordinates for repeat families annotated in RepeatMasker (66) for hg38 using AnnotationHub (“AH111333”). These annotations were used as input to the polyenrich function with the up- or down-regulated DEGs used as the gene set with locusdef=“nearest_gene”. We performed BH correction via p.adjust(method=”fdr”) on resulting p-values to get FDR values and set a significance cutoff of 0.05. All analyses were performed using R (v4.3.1). Plots were generated using EnrichedVolcano (67) and ComplexHeatmap (68, 69) packages.

### Quantitative Telomerase Repeated Amplification (Q-TRAP) Assay

hESC were collected through EDTA-dissociation, centrifuged at 300 × g for 4 min at 4^°^C to pellet. The pellet was washed with PBS. Cells were counted on a Countess (Thermo Fisher) and 500,000 cells per condition were analyzed for telomerase activity using the TRAPeze RT Telomerase Detection Kit (Millipore Sigma. S7710) following manufacturer’s instructions. The cells were lysed in 200 µL of CHAPS lysis buffer containing 200 units/mL of RNase inhibitor (Promega, N2511) on ice for 30 min and then centrifuged at 12000 × g for 20 min at 4^°^C to collect the supernatant. Telomerase activity was then assessed on 5,000 cell equivalents (2 μl of lysate). In each reaction, the telomerase enzyme catalyzed the addition of telomeric repeats to the 3’ end of the oligonucleotide substrate. Following this the extended products are amplified using the *Taq* DNA polymerase (New England BioLabs, M0273S), and TS and fluorescein labelled Amplifluor primers in the prescribed 5X TRAPeze RT reaction mix. The fluorometric signal resulting from the incorporation of the primers into the PCR products during the reaction was analyzed using CFX Opus 96 real-time PCR system (Bio-Rad) and CFX Maestro software. The standard curve for quantification of telomerase activity in individual samples was obtained by serially diluting the control template TSR8 which is an oligonucleotide with identical sequence to the TS primers and 8 telomeric repeats. The control template was diluted in CHAPS lysis buffer to obtain concentrations ranging from 20 amoles/µL to 0.02 amoles/µL and 2 uL of each was used in similar reaction conditions as above obtain the standard curve.

## RESULTS

### APEX nuclease inhibitor impairs TERT transcription in human embryonic stem cells

We previously reported that telomerase-mediated extension of telomeres relied on signaling from DNA repair kinases in an immortalized embryonic neuronal cell line (1). These include the enzymes ataxia telangiectasia mutated (ATM), and ataxia telangiectasia mutated and RAD3-related (ATR). Canonically, DNA repair kinases primarily control the activation of other proteins, but they can also influence the expression of many genes (70). For example, *TERT* gene expression in human ESC was reported to be controlled by poly ADP-ribose polymerase 1 (PARP1) (71), an enzyme central to numerous DNA repair pathways. Based on our previous observations that ATM and ATR control telomerase extension of telomeres (1), we considered the possibility that ATM and ATR may also influence *TERT* gene expression to promote telomere length maintenance. To test the role of ATM and ATR, we employed inhibitors used in our previous study that target the enzymatic activity of ATM and ATR (Ku55933 and Ve-821). As controls we included small molecule inhibitors of DNA protein kinase (DNA-PK, NU-7441) and an inhibitor of APEX1 and APEX2 nuclease function, APEIII which targets APEX nuclease functions. We also included an inhibitor of MEK1 and MEK2 kinases, trametinib, as a positive control, which we recently identified as controlling *TERT* transcription in human stem cells (55, 72). Given the gaps in our knowledge about TERT regulation in normal human stem cells, we chose to study human embryonic stem cells. These cells were cultured under primed stem cell conditions (Fig S1) (73–75). We examined *TERT* mRNA expression by qRT-PCR at exon 2 or exon 14 which are essential for full-length *TERT* transcripts encoding telomerase enzymatic activity. As expected, the positive control for inhibiting *TERT,* MEKi (trametinib, 100 nM, 24 h), suppressed *TERT* transcription by approximately 50% in hESC (Fig. 1A). In contrast, ATMi and ATRi at concentrations and durations that affected affected telomerase assembly (1), showed no effect on *TERT* gene expression (Fig. 1A). These data support our previous model that these kinases primarily impact telomerase assembly and telomere length maintenance at chromosome ends. Unexpectedly, treatment of hESCs with 1 μM APEIII consistently resulted in a 50% reduction in transcription of *TERT* (Fig 1A). This effect was time- and dose-dependent (Fig. 1B, C).

**Figure 1.**
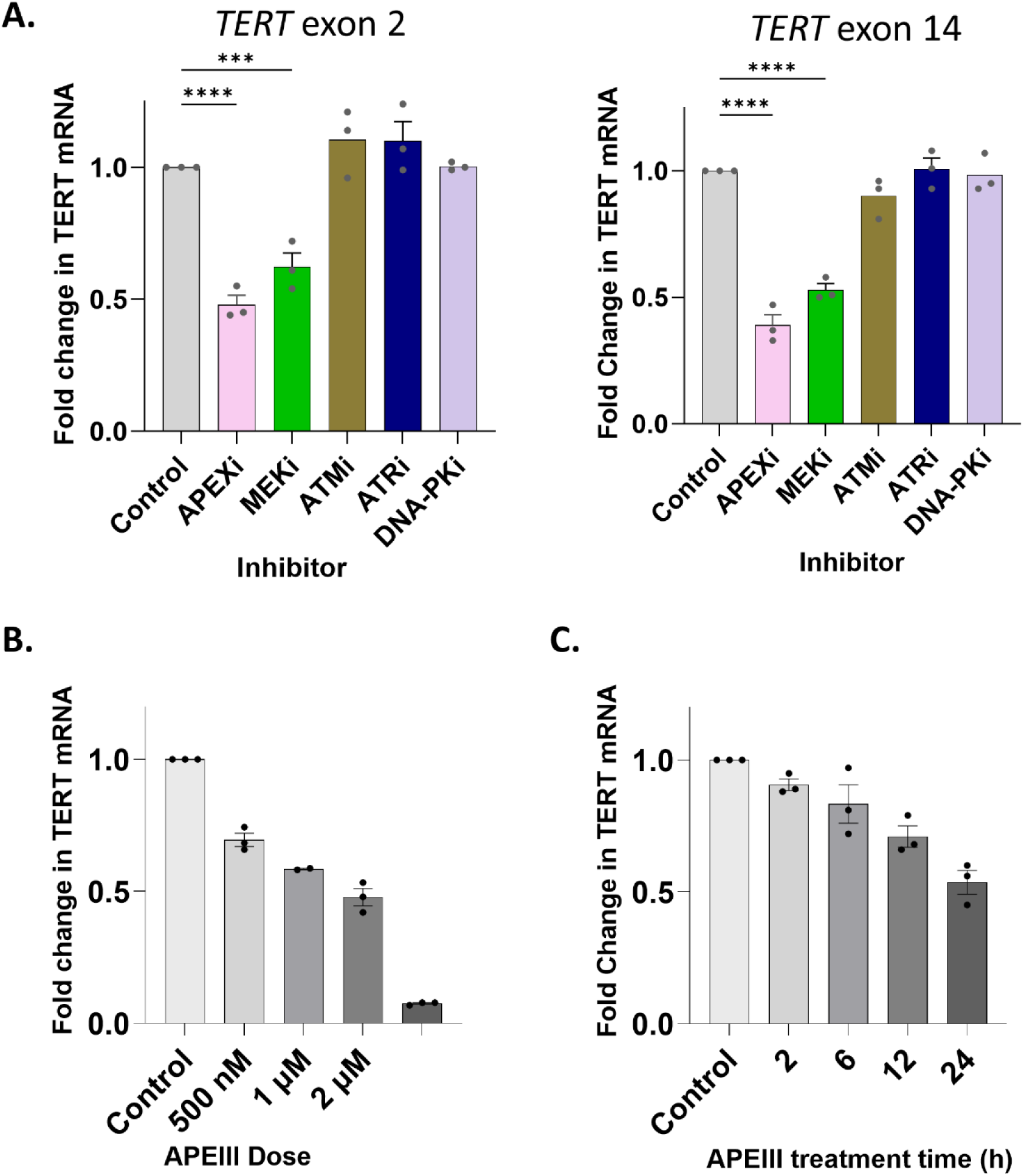
APEX nuclease inhibitor suppresses *TERT* mRNA transcription in human ESC. *TERT* mRNA levels as measured by qRT-PCR vs GAPDH. **(A)** Human embryonic stem cells were treated with the indicated inhibitor for 24 h. APEXi (APEIII, 1 µM), ATMi (KU55933, 5 µM), MEKi (Trametinib, 100 nM), ATRi (VE-821, 400 nM), DNA-PKi (NU-4773, 400 nM).*** P <0.001, One Factor ANOVA, Fisher’s LSD test. **(B)** Human ESC treated with APEIII for 24 h at the indicated dose. **(C}** Cells were treated with APEIII at 1 µM and harvested after the indicated time. Data are mean ±SEM.

### TERT expression in primed hESC is independent of OGG1 and PARP1 activity

The *TERT* proximal promoter is very rich in guanine and cytosine (GC) nucleotides. Its ability to initiate *TERT* transcription is regulated by guanine-rich tertiary structures such as G-quadruplexes that are subject to base oxidation and repair (76, 77). APEX1 and APEX2 play roles in base excision repair (BER). BER employs the 8-oxoguanine N-glycosylase/DNA lyase OGG1 immediately upstream of APEX1 and APEX2 to excise oxidized guanine residues. We therefore hypothesized that APEX2 may promote *TERT* expression in conjunction with OGG1. Intriguingly, the excision of oxidized guanines by OGG1 can promote RAS signaling and gene transcription (78) suggesting this pathways potential to promote *TERT* expression. We first examined the *TERT* promoter for OGG1 occupancy at the *TERT* proximal promoter by ChIP-qPCR but did not observe strong binding (data not shown). However, OGG1 only binds chromatin transiently during the repair of oxidized guanines and is quickly displaced by APEX proteins (79). To test the potential role of OGG1-mediated repair of the *TERT* promoter more fully, we knocked down *OGG1* with siRNA. As PARP1 was reported to play a role in *TERT* expression in hESC, we also included *PARP1* siRNA as a positive control. Knockdown of *OGG1* had little effect on *TERT* expression (Fig. S1A, B). Surprisingly, the knockdown of *PARP1* also had only minor effects on *TERT* expression (Fig S1A, B).

To confirm the presence of oxidized bases in the *TERT* locus and their relationship with *TERT* expression, we modified the Repair Assisted Damage Detection (RADD) assay specifically for oxidative DNA lesions, oxRADD (60). Using a cocktail of FPG and EndoIV to excise oxidized bases and abasic sites, RADD leaves gapped DNA, which is then tagged with a biotinylated dUTP, facilitating the isolation of damaged DNA regions by streptavidin purification for NGS or similar analyses (60). Similarly, a digoxigenin-dUTP can be inserted for fluorescent microscopy analysis using standard immunofluorescence techniques (80).

Using fluorescent microscopy, we first employed the oxRADD cocktail to detect oxidative lesions within single hESC cells. We saw significantly elevated oxidative DNA damage within hESC cells treated with the PARP inhibitor, olaparib (500 nM), for 24 h (Fig S1C). We then employed a modified oxRADD assay on isolated DNA to determine if the *TERT* locus contains oxidative DNA lesions under homeostatic conditions. hECS cells were treated with olaparib (500 nM) for 24 h, preventing the completion of the oxidative repair and allowing naturally occurring oxidized lesions to accumulate. DNA was extracted from these cells and subjected to oxRADD with the insertion of a biotinylated dUTP to allow the isolation of damaged DNA. We then used amplification with specific primers to examine the purified biotinylated DNA for the proximal *TERT* promoter. Increasing DNA damage within the *TERT* promoter will increase the amount of this locus isolated by biotinylation and increase the qPCR signal. As a negative control, we treated a companion sample with the same enzyme cocktail lacking biotin. The treatment of cells with olaparib resulted in elevated amplification of the *TERT* promoter from the oxRADD-processed DNA (Fig. S1D).

As OGG1 relies on PARP1 signaling to initiate repair, we also measured *TERT* mRNA expression in olaparib-treated ESC. We did not observe an effect of PARP1 enzyme inhibition on *TERT* mRNA levels as measured by qRT-PCR of *TERT* exon 2 (Fig. S1E). We interpret these results to indicate that under homeostatic conditions, the *TERT* promoter in hESC sustains oxidative lesions that are repaired in a PARP1-dependent manner. However, while the *TERT* promoter appears to undergo base oxidation and repair in a PARP1-dependent manner in human ESC, it does not affect *TERT* expression, indicating that the effect of APEIII on *TERT* does not involve PARP1 or OGG1.

### APEX2 promotes TERT transcription in human ESC

APEIII was initially characterized as an enzymatic inhibitor of the APEX1 nuclease domain (81) but more recently it was found to phenocopy APEX2 not APEX1 knockdown in a colorectal carcinoma cell line (82). To explore the effect of APEX1 and APEX2 on *TERT* expression we suppressed their expression with siRNA. As negative controls, we included siRNAs targeting two genes, *XRCC6* and *TRIM28*, that have DNA repair functions unrelated to APEX1 and 2. The siRNA against APEX1 reduced *TERT* expression (Fig. 2A), but a stronger effect was observed for the siRNA against APEX2, which consistently lowered *TERT* expression by ∼50%, as measured by levels of exon 2 mRNA (Fig. 2). To enhance our confidence the effects of the siRNA were due to loss of APEX2 protein, we performed immunoblot after siRNA treatment, revealing that APEX2 protein was significantly decreased 72 h following siRNA treatment (Fig 2B, C).

**Figure 2.**
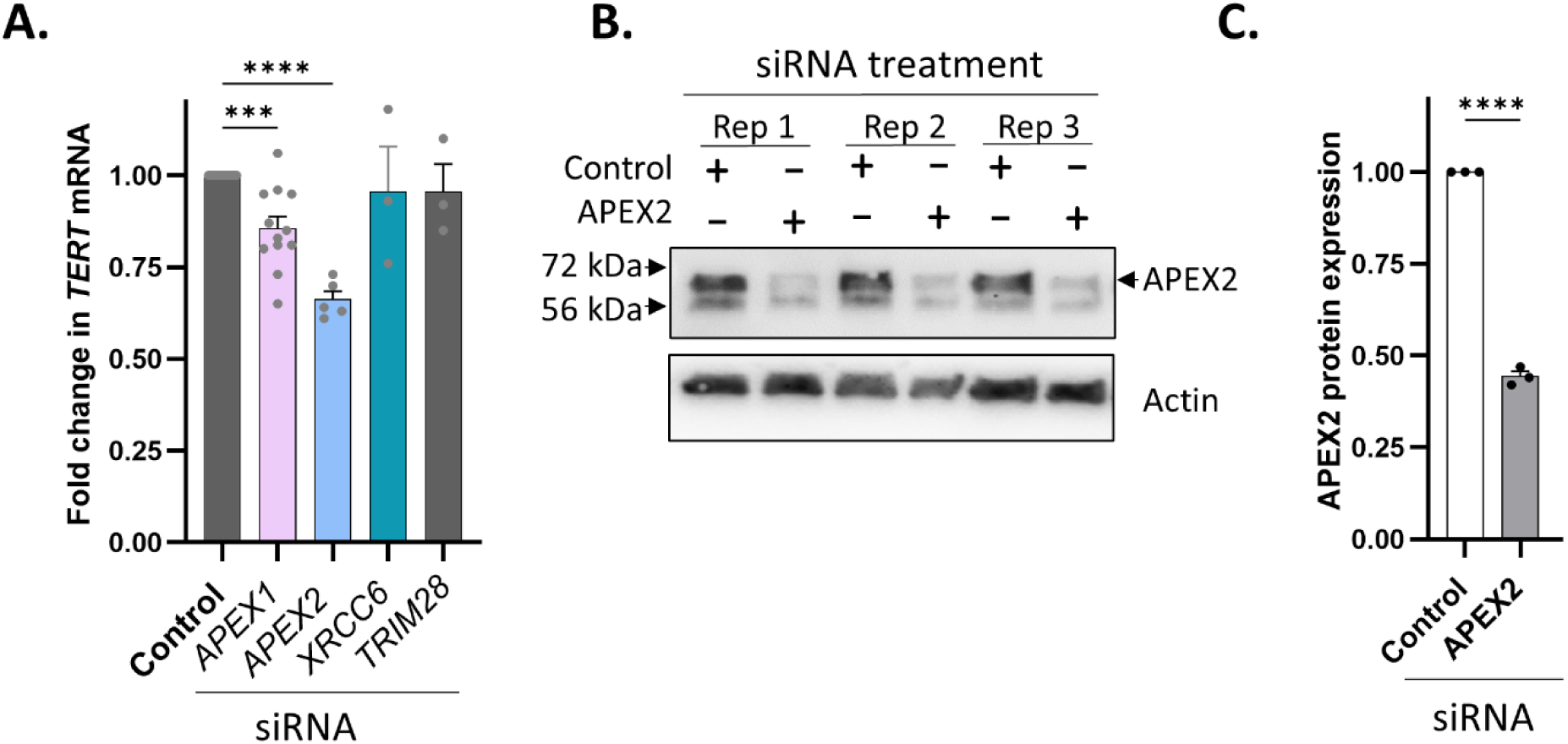
APEX2 knockdown suppresses *TERT* mRNA transcription in human ESC. **(A)** *TERT* mRNA levels as measured by qRT-PCR of exon 2 vs GAPDH. Human embryonic stem cells were treated with the indicated siRNA for 72 h. *** P < 0.01. One factor ANOVA, Fisher’s LSD test. **(B)** lmmunoblots of three independent siRNA knockdowns of APEX2 vs control siRNA in hESC after 72 h. **** P < 0.001, Student’s t-test. **(C)** Quantification of APEX2 knockdowns by immunoblot. Graphs depict mean ±SEM.

### APEX2 knockdown regulates the expression of a subset of genes

Although APEX2 has not been reported to regulate gene expression directly, we hypothesized that transcription of *TERT* was not the only locus impacted by APEX2 loss. An initial small-scale screen of unrelated genes indicated that the effect of APEX2 knockdown on *TERT* was not likely due to widespread suppression of transcription (Fig 3A), as expression of these other genes was unperturbed. To gain insight into how APEX2 influences gene expression, and define the effect on APEX2 gene expression more fully, we performed RNA-seq on triplicate independent ESC samples treated with APEX2 siRNA. These experiments revealed that loss of APEX2 is associated with significant changes in the expression of 347 (FDR < 0.05, Wald test) genes (Fig. 3B, Supplementary Table 3). We performed gene ontology analysis with this set of DEGs and found significant enrichment in pathways associated with translation and gene expression (Figure 3C).

**Figure 3.**
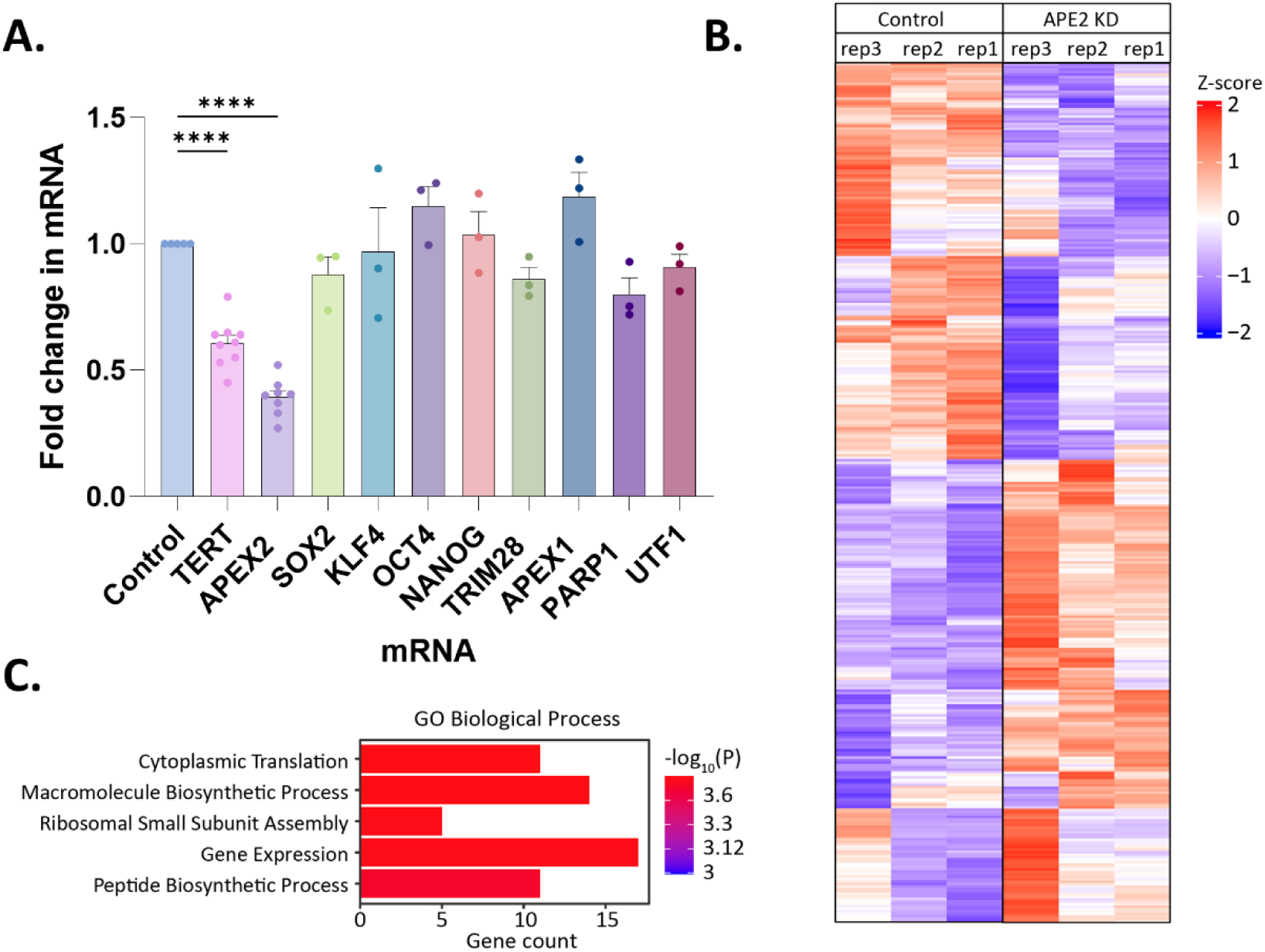
APEX2 impacts mRNA transcription in human ESC. **(A)** mRNA levels of the indicated gene as measured by qRT-PCR. Human embryonic stem cells were treated with APEX2 siRNA for 72 h. **** P< 0.001. One factor ANOVA, Fisher’s LSD test.(B) Heatmap of 347 differentially expressed genes upon APEX2 knockdown in hESC. **(C)** Gene ontology enrichment analysis of differentially expressed genes from (B). See also Supplementary Table 3.

### APEX2 regulation of TERT is cell-type dependent

To assess the role of APEX2 on *TERT* expression in other immortalized cells, we tested APEX2 siRNA in telomerase-positive melanoma (A101) and glioblastoma stem cells (GSC75). Loss of APEX2 in melanoma cells impacted *TERT* expression to a similar degree observed in hESC (Fig. 4A). Surprisingly, APEX2 knockdown had little impact on *TERT* expression in GSC75 cells (Fig. 4A) indicating that the effect of this factor in promoting *TERT* expression is cell-type specific. To determine whether other pluripotent stem cells rely on APEX2 for *TERT* expression, we tested induced pluripotent stem cells (iPSC) derived from CD34^+^ bone marrow cells. Despite efficient knockdown of APEX2, similar to GSC and melanoma, *TERT* expression was unaffected in iPSC (Fig 4B).

**Figure 4.**
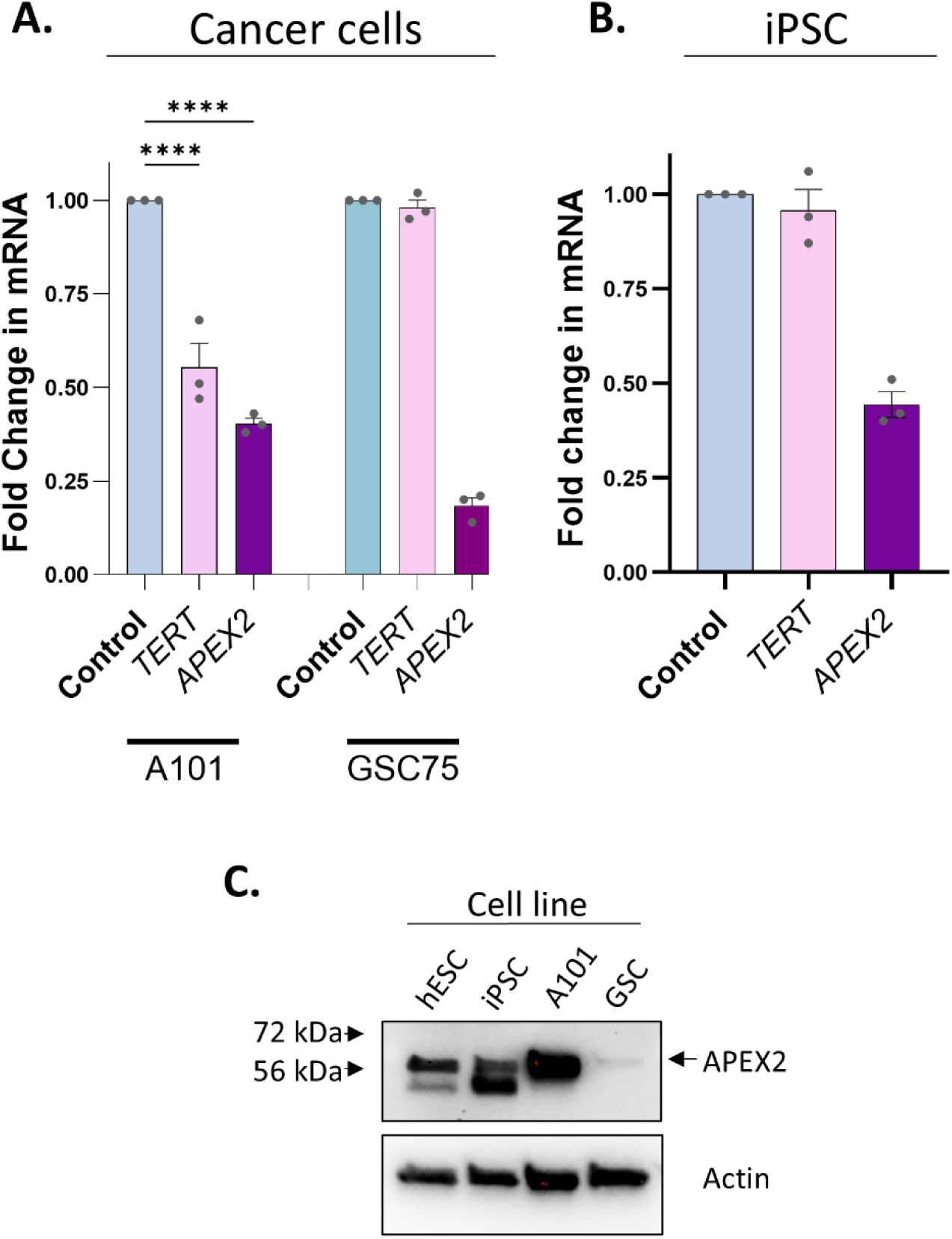
Cell-type dependent effects of APEX2 on *TERT* expression. *TERT* mRNA levels as measured by qRT-PCR of exon 2. **(A)** Human melanoma cell line Al0l orglioma stem cell lines GSC75 and GSC39 treated with 50 pmol APEX2 siRNA for 72 h. **(B)** hlPSC treated with 50 pmol APEX2 siRNA for 72 h. *** P <0.01, One Factor ANOVA, Fisher’s LSD test. **(C}** lmmunoblots for APEX2 and beta actin protein expression in hESC, iPSC, melanoma cells Al0l and glioblastoma cells GSC75.

The cell-specificity of APEX2 knockdown on *TERT* expression suggested differences in APEX2 function between cell types. We therefore assessed the levels of APEX2 in hESC, iPSC, A101, and GSC75. The two cell types that displayed APEX2 independence for *TERT* expression, GSC75 and iPSC, had the lowest levels of APEX2 protein relative to actin (Fig 4C). Together, this suggests that APEX2 expression levels may play a role in the dependence of *TERT* levels on this protein.

### APEX2 displays elevated binding near a TERT tandem repeat

How APEX2 promotes efficient gene expression is unclear. APEX2 lacks homology to the APEX1 redox effector domain that mediates the ability of APEX1 to promote transcription. We considered the possibility that APEX2 promoted efficient *TERT* transcription by binding to the *TERT* proximal promoter, as this region is important for regulating *TERT* in stem cells. However, ChIP-qPCR experiments indicated relatively low levels of binding in this region (Fig. 5).

**Figure 5.**
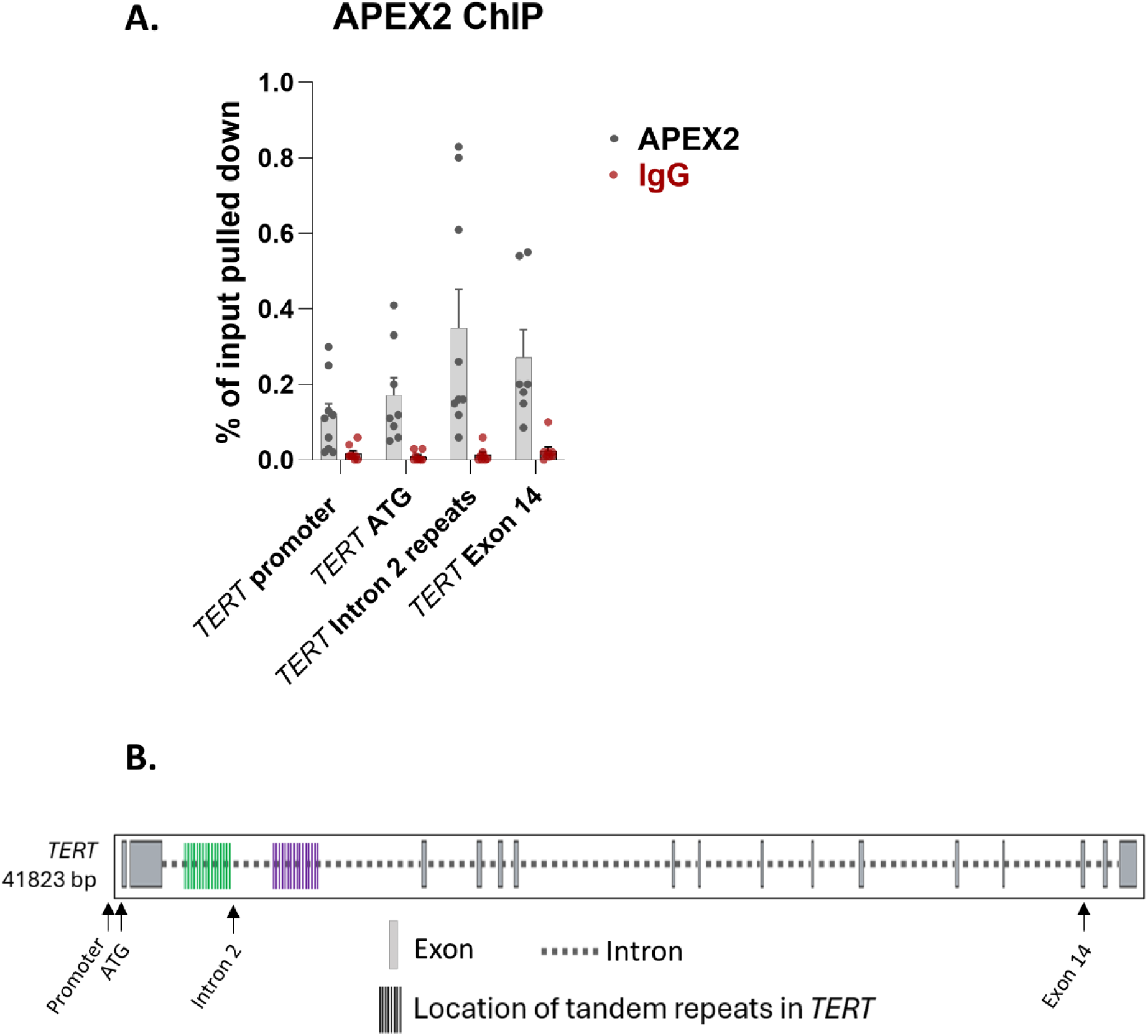
APEX2 binding is highest in the *TERT* gene body in human ESC. **(A)** Chromatin immunoprecipitation of human ESC. Pull down % is calculated vs input DNA. Data are means ±SEM. **(B)** Primer positions in the *TERT* gene for detection of APEX2 binding are indicated by black arrows.

DNA damage in repetitive sequences located in genes can interfere with efficient mRNA expression (45–51, 83). These include GC-rich sequences as well as mammalian wide interspersed repeats (MIR) and transposable elements such as long interspersed nuclear elements (LINE), short interspersed nuclear elements (SINE), and long terminal repeat retrotransposons (LTR) (84). The *TERT* locus contains repetitive DNA features that pose elevated risks for DNA damage (51, 84). The *TERT* MIRs comprise minisatellite DNA with a variable number of tandem repeats (VNTR) such as those in intron 2 (51, 84). These repetitive elements have been implicated in regulating the expression of *TERT* in artificial constructs (85). We posited that *TERT* may rely on APEX2 to frequently repair these regions to minimize their ability to interrupt transcription. Specifically, we hypothesized that *TERT* recruits APEX2 to these sequences and tested this by ChIP for APEX2 in hESC. Quantitative PCR primers were designed to amplify a region approximately 90 bp from the 3’ boundary VNTR in *TERT* intron 2. We also tested APEX2 occupancy at the *TERT* transcriptional start site and at exon 14 near the 3’ of the gene. We observed the highest APEX2 binding adjacent to the tandem repeats in *TERT* intron 2 (Fig. 5) suggesting that APEX2 may act locally at these repetitive sequences to promote efficient expression.

### Expression of genes with MIR elements are impaired by APEX2 silencing

Based on these observations at *TERT* we assessed if other genes impacted by APEX2 knockdown were enriched for similar repetitive elements. We determined whether APEX2-regulated genes from our RNA-seq analysis contained specific types of repeat elements within the gene body. We utilized the Poly-Enrich method (65) to calculate enrichments of major repeat families annotated in the RepeatMasker database (66) within the gene bodies of up- or down-regulated DEGs. These analyses revealed that down-regulated genes that rely on APEX2 for efficient expression were significantly enriched for Mammalian-wide interspersed repeats (MIR; FDR = 0.04; 144 genes) and L2 repeats (FDR = 0.03; 147 genes) (Fig. 6A, B, Supplementary Table 4). In contrast, down-regulated genes were depleted for Alu and ERV1 repeat families. Upregulated genes consistently showed the opposite effect for each repeat family. These results suggest that APEX2 is required for efficient transcription of a subset of genes such as *TERT* with specific types of DNA repeats.

**Figure 6.**
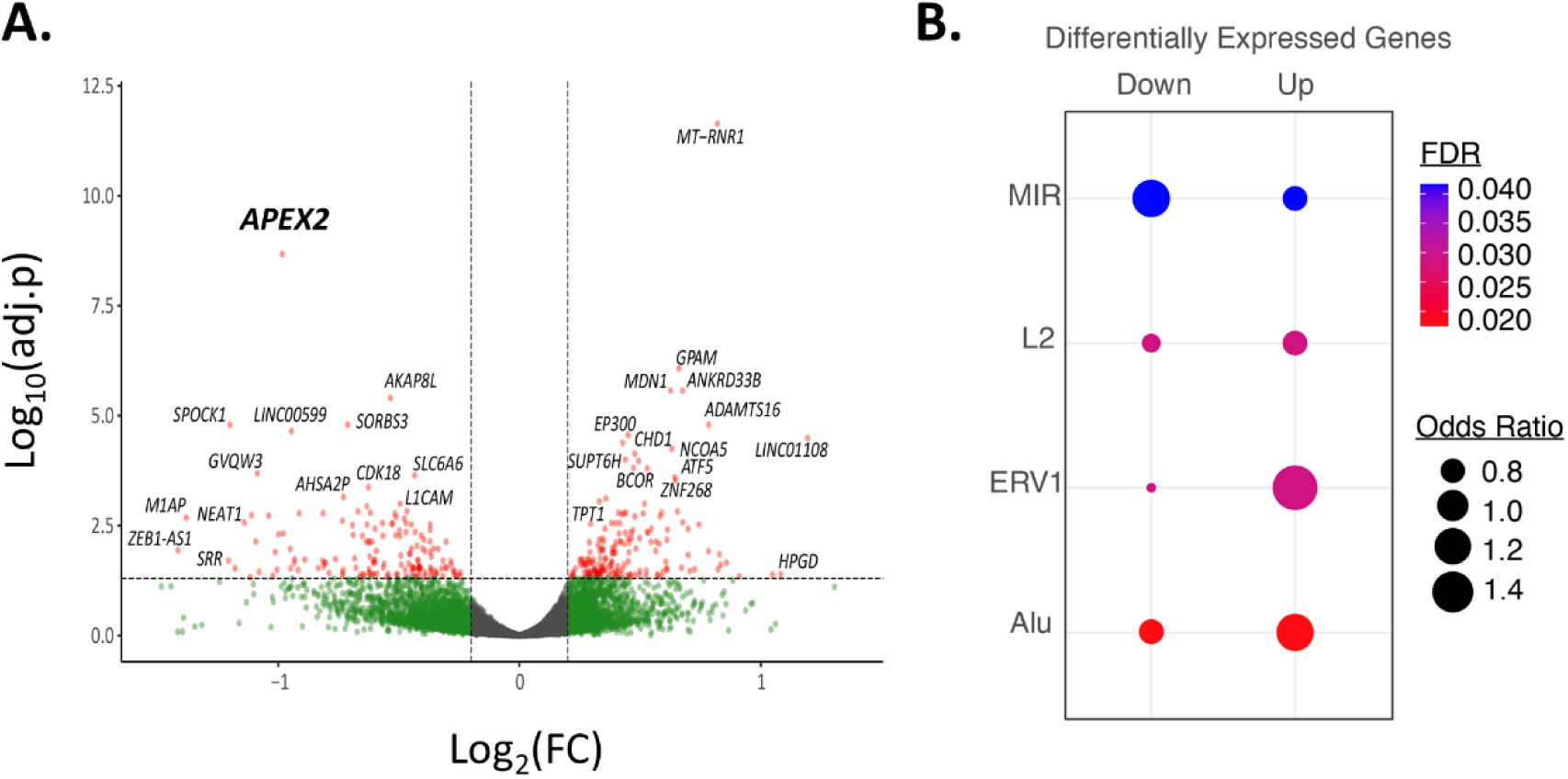
APEX2 impacts mRNA transcription of genes enriched for specific types of repetitive sequences. **(A)** Volcano plot of RNA-seq results after APEX2 knockdown. Significant DEGs in red denoting FDR<0.05 and log2FC 2: 0.5. **(B)** Enrichment of up- and down-regulated DEGs for specific families of repetitive elements. See also Supplementary Table 4.

### APEX2 promotes telomerase enzyme activity in ESC

To determine if the effect of inhibiting APEX2 translated into decreased telomerase enzyme activity we repeated the APEX2 siRNA knockdown and APEIII inhibitor treatment and analyzed the cells by quantitative telomere repeat amplification protocol (Q-TRAP)(86). Treatment of cells with siRNA against APEX2 consistently reduced telomerase enzyme activity (Fig. 7A, S2), while knockdown of APEX1 did not have this effect (data not shown). Similarly, inhibition of APEX nuclease activity also impaired telomerase activity (Fig. 7B). The impact of these treatments on telomerase activity was comparable to the level of *TERT* transcriptional suppression by these treatments (Fig. 1, 2). We conclude that APEX2 is required for efficient telomerase expression in hESC and is likely important for telomere length maintenance during embryogenesis.

**Figure 7.**
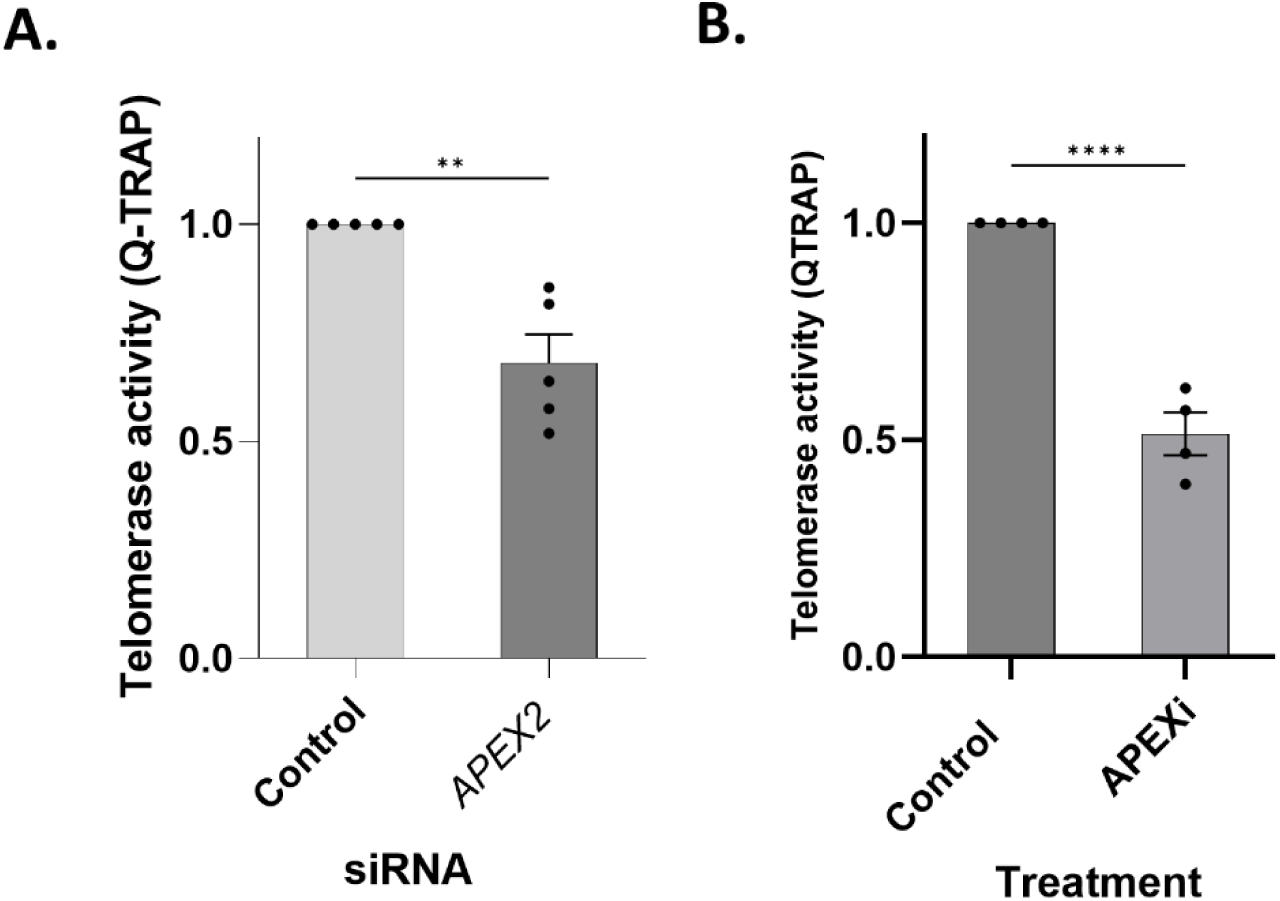
APEX2 inhibition suppresses telomerase enzyme activity in hESC. Telomerase activity determined by quantitative TRAP assay in hESC. **(A)** Cells treated with 50 pmol APE2 siRNA pool vs control siRNA for 72 hours. **(B)** Cells treated with 1 µM APEIII for 24 hours. Graph depicts mean± SEM, n=3. ** P < 0.01, Student’s t-test.

## DISCUSSION

The integration between telomere maintenance and DNA repair proteins is extensive and highly conserved across kingdoms. For example, we previously showed that ATM and ATR are critical for telomere maintenance in cancer cells (1). Telomerase reverse transcriptase is a specialized arm of the cellular DNA repair mechanisms that counteracts telomere erosion, critically sustaining the perpetual growth of pluripotent stem cells. We tested the contribution of a number of DNA repair proteins to the expression of *TERT* and found that APEX2 is required for efficient expression of a subset of genes enriched for specific DNA features, including *TERT*. We also show that hESC relied on APEX2 to maintain full telomerase enzyme activity. APEX2 was recently reported to be important for telomere length maintenance by DNA double-strand break repair (87) indicating that APEX2 likely plays multiple roles in promoting telomere length maintenance.

How does APEX2 promote efficient gene expression? Our RNA-seq analysis suggests that genes impacted by APEX2 loss are enriched for specific types of repetitive DNA sequences. These repetitive elements are predicted to undergo elevated levels of DNA damage during replication and transcription. Failure to repair such intragenic DNA damage can impair gene transcription by interfering with RNA polymerase passage (51). Given the enrichment of these DNA repetitive elements in the APEX2 gene set, one possibility is that APEX2 may be important for efficiently repairing these sequences to promote RNA polymerase II-mediated transcription. Our observation of elevated recruitment of APEX2 adjacent to the VNTR sequences in *TERT* intron 2 supports this hypothesis. Future studies examining the recruitment of APEX2 to repetitive sequences in other genes may provide further insight into how APEX2 influences the expression of these genes.

Despite high levels of homology, APEX2 differs structurally and functionally from APEX1. APEX2 harbors a single-stranded DNA (ssDNA)-binding Zf-GRF domain as well as a Proliferating Cell Nuclear Antigen-interacting motif (PIP) at its C terminus (88). Additional differences include APEX2’s weaker endonuclease activity and increased 3′- to 5′-exonuclease and 3′-phosphodiesterase action (44, 89). Intriguingly, APEX2-deficient mice show defects in certain bone marrow progenitor cells after chemotherapy (90), an effect dependent on p53 (91, 92). Although the precise mechanisms by which APEX2 supports these cells is unknown, our study suggests that the ability of APEX2 to promote efficient expression of a subset of genes may contribute to this biological role. An interesting observation in our study is the cell type-specific effect of APEX2 on *TERT* expression. While our study did not determine the basis for this, our data suggest that APEX2 expression levels may play a role and warrant further tests in different cell types and in additional donor cells and their reliance on APEX2 to promote gene expression.

Few prior studies have identified specific regulators of *TERT* expression in validated models of normal human stem cells (93, 94). Two such studies reported that proteins involved in DNA repair, PARP1 and KLF4 controlled *TERT* gene expression in human ESC (71, 95). We did not find evidence that PARP1 was a major factor in promoting efficient *TERT* gene expression under the growth conditions we used. Since ESC gene expression can be influenced by the growth conditions (74, 75), one possible explanation is the different media used in our study (Essential 8 Flex media). Studies of *TERT* by Teng and colleagues maintained hESC with murine feeder cells, which may have resulted in different gene regulation (71, 95). The growth conditions used in our study maintained the cells with phenotypes consistent with primed ESCs (96). Primed hESC, including those in our study, lack substantial expression of KLF4, while naïve hESC cells express KLF4. Thus, an intriguing possibility is that regulation of *TERT* gene expression may undergo dynamic changes during the transition from naive to primed developmental stages. Future studies will be required to determine if *TERT* undergoes changes in transcriptional regulation from primed to naïve states.

*TERT* expression is driven by RAS signaling in many cell types including hESC (96) and RAS signaling was recently reported to promote telomere lengthening in animal models (97). Intriguingly, induction of RAS signaling was observed following release of 8-oxo-dG during OGG1-mediated base excision repair (78), suggesting the potential for this pathway to promote *TERT* expression. However, several lines of evidence in our study (Fig. S1A-E) suggest that the repair of oxidized guanines by PARP1/OGG1 does not participate in the APEX2-mediated *TERT* expression we report here.

Telomerase is suppressed after embryogenesis in most tissues to limit the unscheduled proliferation of normal somatic cells. The regulation of *TERT* transcripts primarily mediates this control. Understanding the regulatory control of the *TERT* gene locus in human stem cells remains limited in part due to the dominance of murine stem cell models in the field, which display different telomerase regulation and telomere dynamics. Our study identifies new roles for the DNA repair protein APEX2 as well as a new cell type-specific mechanism controlling the expression of *TERT* in a subset of human stem and cancer cells.

**Table 2.** List of antibodies.

| Antibodies for western blot | Company | Identifier |
| --- | --- | --- |
| APEX2 | Cell Signaling Technology | 74728S |
| Actin | Santa Cruz Technology | sc-47778 |
| **Antibodies for ChIP** |  |  |
| IgG | Cell Signaling Technology | 2729S |
| APEX2 | Cell Signaling Technology | 74728S |
| APEX2 | Genetex | GTX80642 |

**Table 1.**
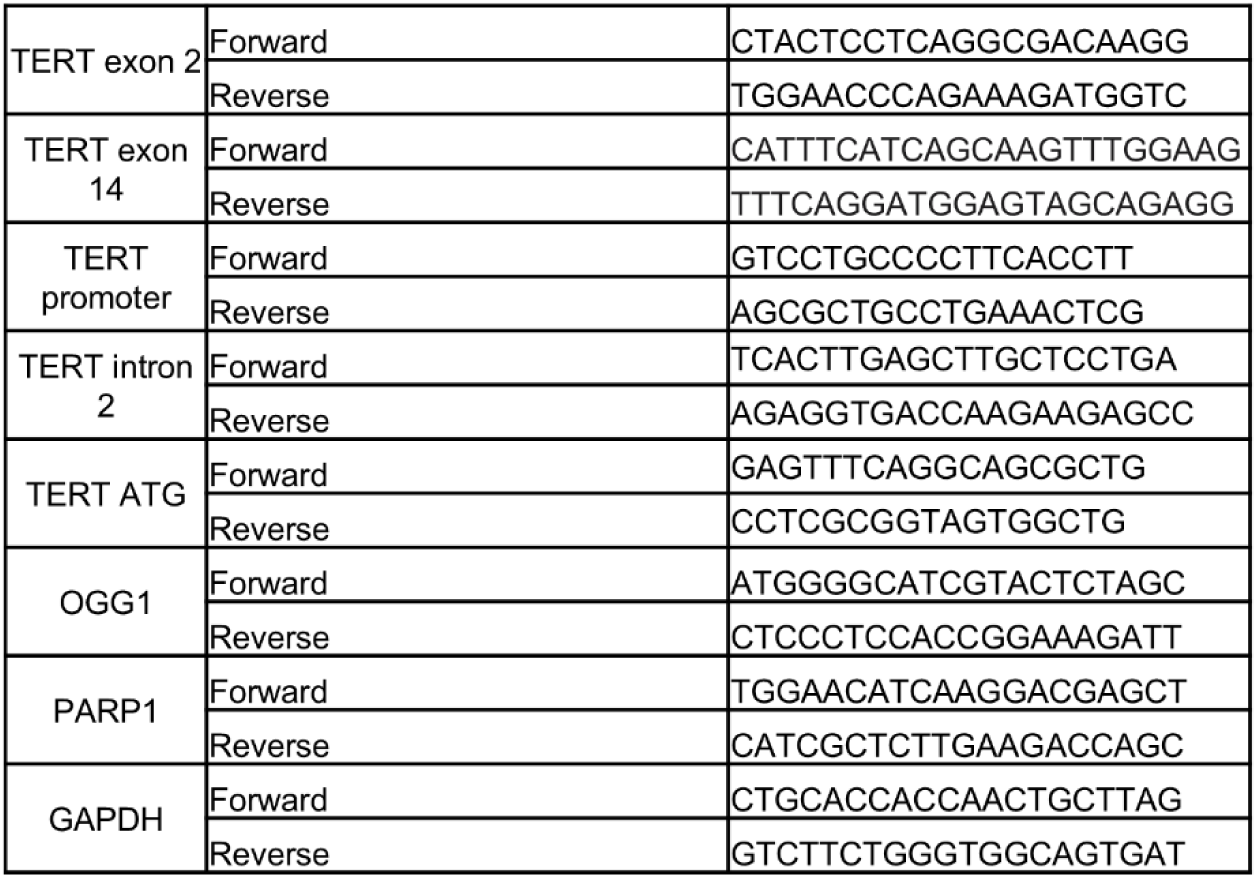
List of primers.

## Supplementary Information

**Figure S1.**
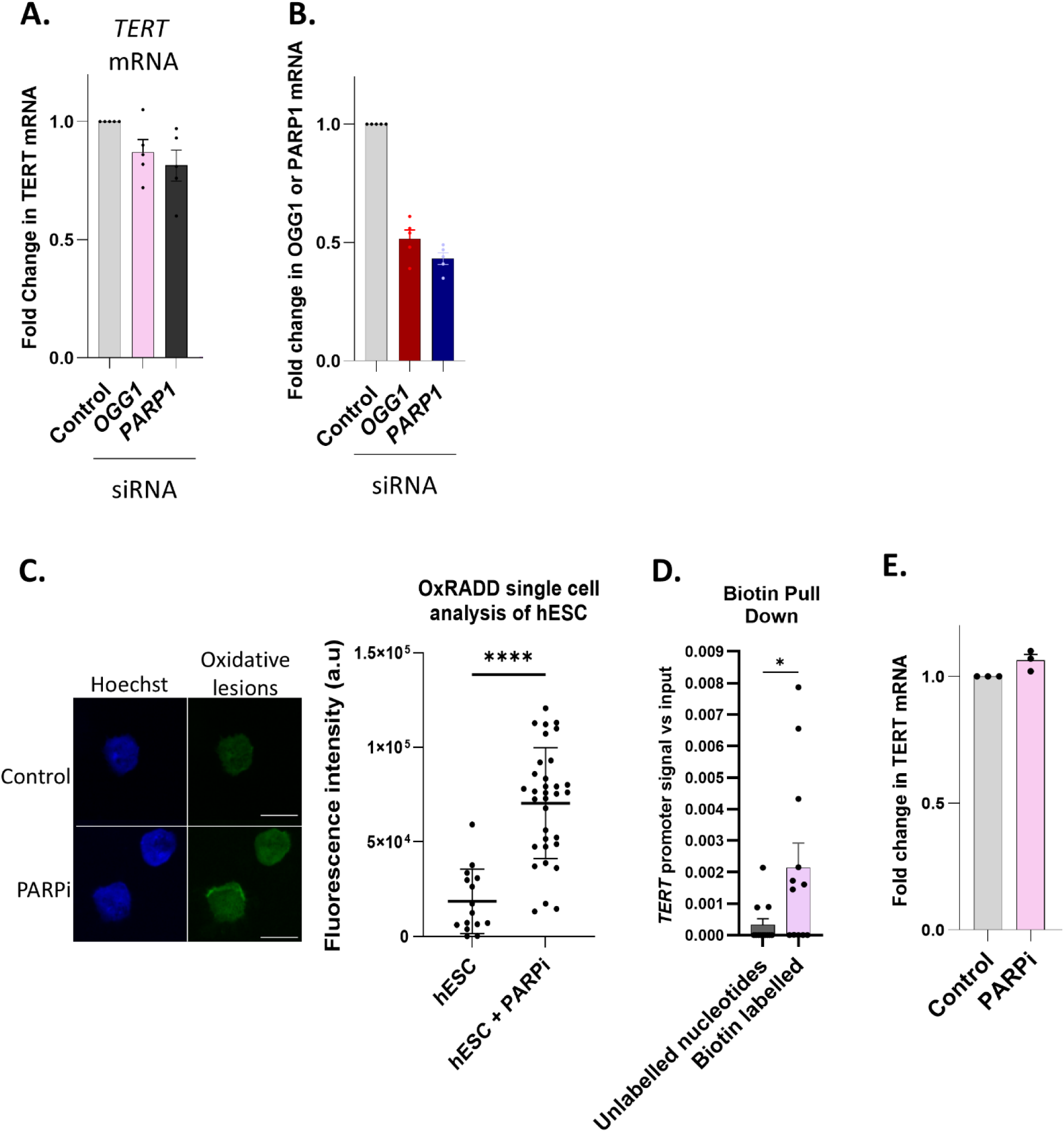
Base oxidation in the *TERT* promoter detected by DNA lesion repair reaction in hESC. **(A), (B)** hESC were treated with siRNA against OGGl or PARPl for 72 h. Expression of the indicated mRNA was measured by qRT-PCR vs GAPDH. Data are +/-SEM for n=3. **(C)** Detection of oxidized bases in hESC treated with olaparib (500 nM, 24 h). Left images are a montage of the cells used for analysis. Scale bar 25 µm. Right images are the fluorescent signal quantified for the oxidative lesions. Significance was calculated using a Welch’s t-test. **(D)** Biotin-pull down and of labelled oxidized bases detected by qPCR quantification of *TERT* promoter. **(E)** *TERT* mRNA levels as measured by qRT-PCR of exon 2 in hESC treated with olaparib (500 nM, 24 h). Data are ±SEM for n=3.

**Figure S2.**
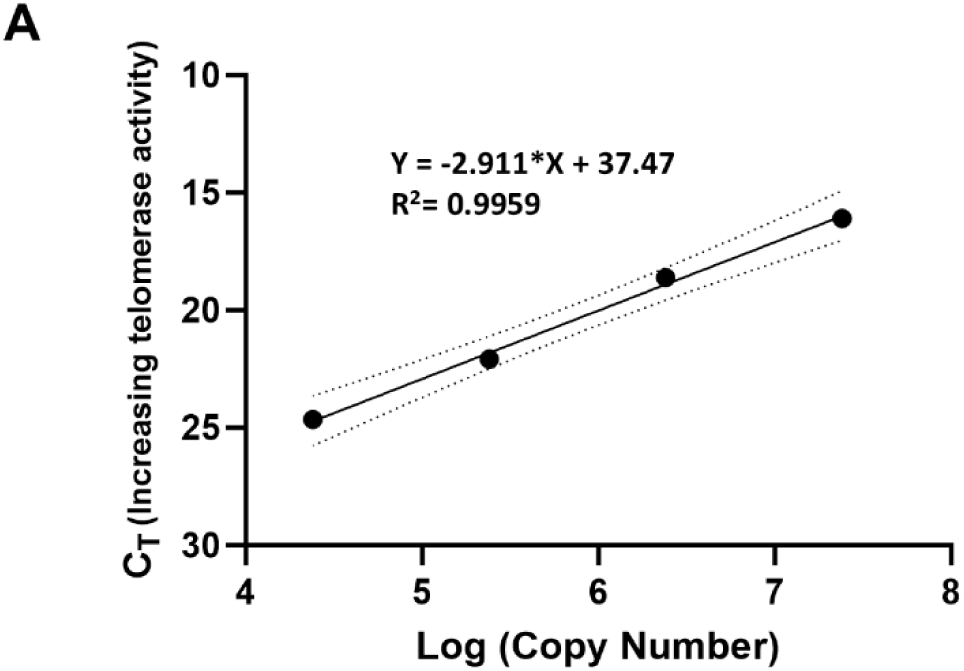
Logarithmic plot of the TSR8 standard curve for quantification of telomerase activity. Graph depicts the CT values corresponding fluorometric signal obtained on the amplification of different dilutions of TSR8 template. Data are +/-SEM for n=3.

